# Sybodies targeting the SARS-CoV-2 receptor-binding domain

**DOI:** 10.1101/2020.04.16.045419

**Authors:** Justin D. Walter, Cedric A.J. Hutter, Iwan Zimmermann, Marianne Wyss, Jennifer Earp, Pascal Egloff, Michèle Sorgenfrei, Lea M. Hürlimann, Imre Gonda, Gianmarco Meier, Sille Remm, Sujani Thavarasah, Philippe Plattet, Markus A. Seeger

## Abstract

The COVID-19 pandemic, caused by the novel coronavirus SARS-CoV-2, has resulted in a global health and economic crisis of unprecedented scale. The high transmissibility of SARS-CoV-2, combined with a lack of population immunity and prevalence of severe clinical outcomes, urges the rapid development of effective therapeutic countermeasures. Here, we report the generation of synthetic nanobodies, known as sybodies, against the receptor-binding domain (RBD) of SARS-CoV-2. In an expeditious process taking only twelve working days, sybodies were selected entirely *in vitro* from three large combinatorial libraries, using ribosome and phage display. We obtained six strongly enriched sybody pools against the isolated RBD and identified 63 unique anti-RBD sybodies which also interact in the context of the full-length SARS-CoV-2 spike ectodomain. Among the selected sybodies, six were found to bind to the viral spike with double-digit nanomolar affinity, and five of these also showed substantial inhibition of RBD interaction with human angiotensin-converting enzyme 2 (ACE2). Additionally, we identified a pair of anti-RBD sybodies that can simultaneously bind to the RBD. It is anticipated that compact binders such as these sybodies could feasibly be developed into an inhalable drug that can be used as a convenient prophylaxis against COVID-19. Moreover, generation of polyvalent antivirals, via fusion of anti-RBD sybodies to additional small binders recognizing secondary epitopes, could enhance the therapeutic potential and guard against escape mutants. We present full sequence information and detailed protocols for the identified sybodies, as a freely accessible resource.

## INTRODUCTION

The ongoing pandemic arising from the emergence of the 2019 novel coronavirus, SARS-CoV-2, demands urgent development of effective antiviral therapeutics. Several factors contribute to the adverse nature of SARS-CoV-2 from a global health perspective, including the absence of herd immunity [1], high transmissibility [2, 3], the prospect of asymptomatic carriers [4], and a high rate of clinically severe outcomes [5]. Moreover, a vaccine against SARS-CoV-2 is unlikely to be available for at least 12-18 months [6], despite earnest development efforts [7, 8], making alternative intervention strategies paramount. In addition to offering relief for patients suffering from the resulting COVID-19 disease, therapeutics may also reduce the viral transmission rate by being administered to asymptomatic individuals subsequent to probable exposure [9]. Finally, given that SARS-CoV-2 represents the third global coronavirus outbreak in the past 20 years [10, 11], development of rapid therapeutic strategies during the current crises could offer greater preparedness for future pandemics.

Akin to all coronaviruses, the viral envelope of SARS-CoV-2 harbors protruding, club-like, multidomain spike proteins that provide the machinery enabling entry into human cells [12–14]. The spike ectodomain is segregated into two regions, termed S1 and S2. The outer S1 subunit of SARS-CoV-2 is responsible for host recognition via interaction between its C-terminal receptor-binding domain (RBD) and human angiotensin converting enzyme 2 (ACE2), present on the exterior surface of airway cells [14, 15]. While there is no known host-recognition role for the S1 N-terminal domain (NTD) of SARS-CoV-2, it is notable that S1 NTDs of other coronaviruses have been shown to bind host surface glycans [12, 16]. In contrast to spike region S1, the S2 subunit contains the membrane fusion apparatus, and also mediates trimerization of the ectodomain [12–14]. Prior to host recognition, spike proteins exist in a metastable pre-fusion state wherein the S1 subunits lay atop the S2 region and the RBD oscillates between “up” and “down” conformations that are, respectively, accessible and inaccessible to receptor binding [12, 17, 18]. Upon processing at the S1/S2 and S2’ cleavage sites by host proteases as well as engagement to the receptor, the S2 subunit undergoes dramatic conformational changes from the pre-fusion to the post-fusion state. Such structural rearrangements are associated with the merging of the viral envelope with host membranes, thereby allowing injection of the genetic information into the cytoplasm of the host cell [19, 20].

Coronavirus spike proteins are highly immunogenic [21], and several experimental approaches have sought to target this molecular feature for the purpose of viral neutralization [22]. The high specificity, potency, and modular nature of antibody-based antiviral therapeutics has shown exceptional promise [23–25], and the isolated, purified RBD has been a popular target for the development of anti-spike antibodies against pathogenic coronaviruses [26–29]. However, binders against the isolated RBD may not effectively engage the aforementioned pre-fusion conformation of the full spike, which could account for the poor neutralization ability of recently described single-domain antibodies that were raised against the RBD of SARS-CoV-2 [30]. Therefore, to better identify molecules with qualities befitting a drug-like candidate, it would be advantageous to validate RBD-specific binders in the context of the full, stabilized, pre-fusion spike assembly [13, 31].

Single domain antibodies based on the variable VHH domain of heavy-chain-only antibodies of camelids – generally known as nanobodies – have emerged as a broadly utilized and highly successful antibody fragment format [32]. Nanobodies are small (12-15 kDa), stable, and inexpensive to produce in bacteria and yeast [33], yet they bind targets in a similar affinity range as conventional antibodies. Due to their minimal size, they are particularly suited to reach hidden epitopes such as crevices of target proteins [34]. We recently designed three libraries of synthetic nanobodies, termed sybodies, based on elucidated structures of nanobody-target complexes (Fig. 1A) [35, 36]. Sybodies can be selected against any target protein within twelve working days, which is considerably faster than natural nanobodies, which requires the repetitive immunization during a period of two months prior to binder selection by phage display [36] (Fig. 1C). A considerable advantage of our platform is that sybody selections are carried out under defined conditions — in case of coronavirus spike proteins, this offers the opportunity to generate binders recognizing the metastable pre-fusion conformation [13, 14]. Finally, due to the feasibility of inhaled therapeutic nanobody formulations [37], virus-neutralizing sybodies could offer a convenient and direct means of prophylaxis.

**Figure 1.**
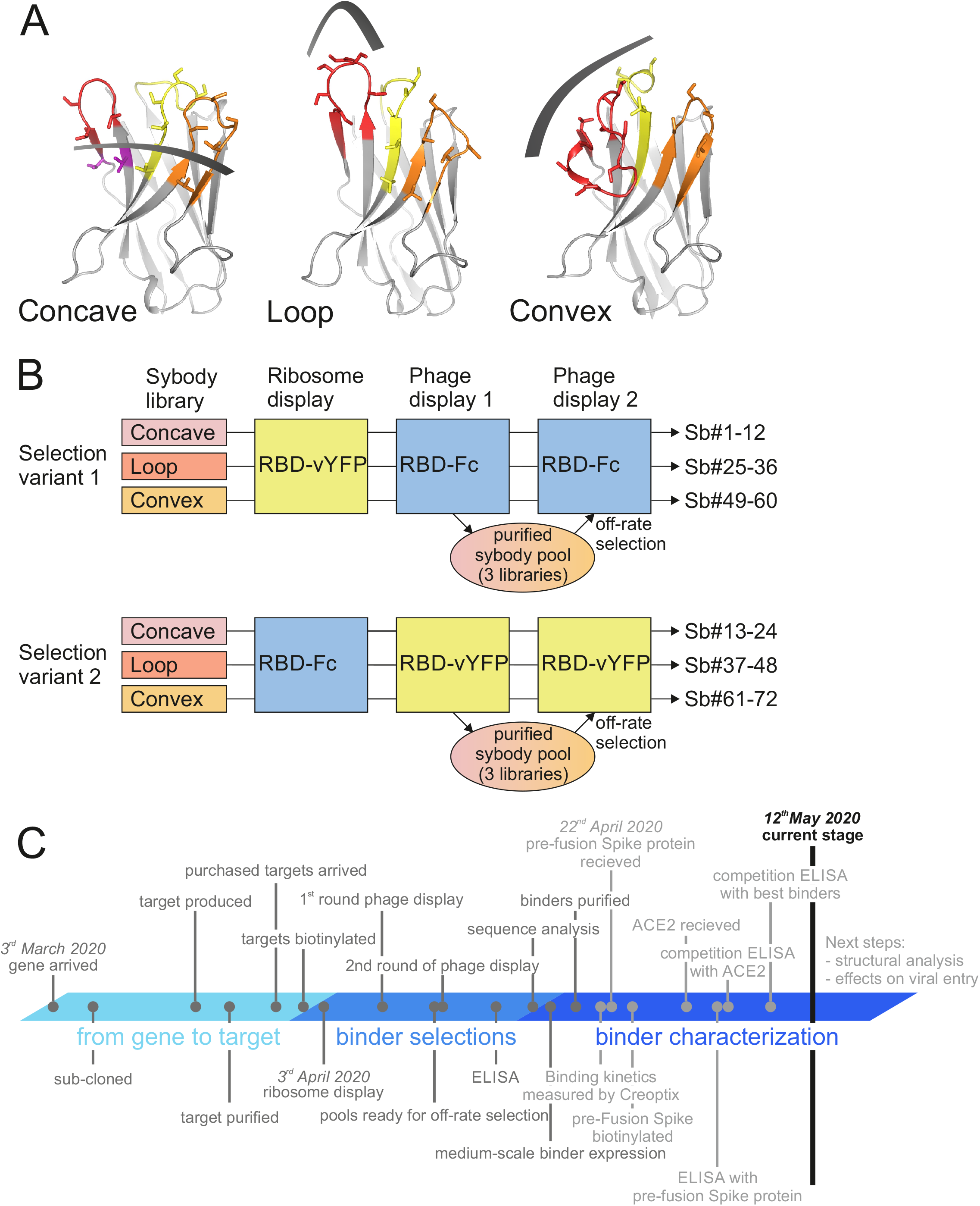
Sybody selections against SARS-CoV-2 RBDs. (**A**) Randomized surface of the three sybody libraries concave, loop and convex. CDR1 in yellow, CDR2 in orange, CDR3 in red. Randomized residues are depicted as sticks. (**B**) Selection scheme. A total of six independent selection reactions were carried out, including a target swap between ribosome display and phage display round. Enriched sybodies of phage display round 1 of all three libraries were expressed and purified as a pool and used to perform an off-rate selection in phage display round 2. (**C**) Time line of this sybody selection process. Please note that this is an intermediate report.

Here, we report of *in vitro* selection and characterization of sybodies against the RBD of SARS-CoV-2 spike protein. Two independently prepared RBD constructs were used for *in vitro* sybody selections, and resulting single clones that could bind the full spike ectodomain were sequenced, expressed, and purified. Six unique sybodies show favorable binding affinity to the SARS-CoV-2 spike, and five of these were also found to substantially attenuate the interaction between the viral RBD and human ACE2. Moreover, pairs of sybodies were identified that can simultaneously bind to the RBD. We present all sequences for these clones, along with detailed protocols to enable the community to freely produce and further characterize these SARS-CoV-2 binders.

## RESULTS AND DISCUSSION

### Purification and biotinylation of target proteins

Based on sequence alignments with isolated RBD variants from SARS-CoV-1 that were amenable to purification and crystallization [29, 38], a SARS-CoV-2 RBD construct was designed, consisting of residues Pro330—Gly526 fused to Venus YFP (RBD-vYFP). This construct was expressed and secreted from Expi293 cells, and RBD-vYFP was extracted directly from culture medium supernatant using an immobilized anti-GFP nanobody [39], affording a highly purified product with negligible background contamination. Initial efforts to cleave the C-terminal vYFP fusion partner with 3C protease resulted in unstable RBD, so experiments were continued with full RBD-vYFP fusion protein. To account for the presence of the vYFP fusion partner, a second RBD construct, consisting of a fusion to murine IgG1 Fc domain (RBD-Fc), was commercially acquired. To remove any trace amines, buffers were exchanged to PBS via extensive dialysis. Proteins were chemically biotinylated, and the degree of biotinylation was assessed by a streptavidin gel-shift assay and found to be greater than 90 % of the target proteins [40]. We note that while both RBD fusion proteins were well-behaved, a commercially acquired purified full-length SARS-CoV-2 spike ectodomain construct (ECD) was found to be aggregation-prone. Very recently, we also produced an engineered spike protein ectodomain containing two point mutations known to stabilize the pre-fusion state, an inactivated furin cleavage site, and a C-terminal trimerization motif [13, 14, 31]. While this purified pre-fusion spike (PFS) had not yet been available for binder selections and characterization by grating-coupled interferometry, it was used to conduct ELISAs in order to identify selected sybodies which recognize the RBD in the pre-fusion context (see below).

### Sybody selections

Since both our RBD constructs bear additional fusion domains (Fc of mouse IgG1 and vYFP, respectively), sybody selections were carried out with a “target swap” approach (Fig. 1B). Hence, selections with the three sybody libraries (concave, loop and convex) were started with the RBD-vYFP construct using ribosome display, and the RBD-Fc construct was then used for the two phage display rounds (selection variant 1: RBD-vYFP/RBD-Fc/RBD-Fc) and *vice versa* (selection variant 2: RBD-Fc/RBD-vYFP/RBD-vYFP). Accordingly, there were a total of six selection reactions (Table 1, Fig. 1B). To increase the average affinity of the isolated sybodies, we included an off-rate selection step using the pre-enriched purified sybody pool after phage display round 1 as competitor. To this end, sybody pools of all three libraries of the same selection variant were sub-cloned from the phage display vectors into the sybody expression vector pSb_init. Subsequently, the two separate pools (all sybodies of selection variants 1 and 2, respectively) were expressed and purified. The purified pools were then added to the panning reactions of the respective selection variant in the second phage display round. Thereby, re-binding of sybody-phage complexes with fast off-rates was suppressed. Enrichment of sybodies against the RBD was monitored by qPCR. Already in the first phage display round, the concave and loop sybodies of selection variant 2 showed enrichment factors of 7 and 3, respectively (Table 1). After the second phage display round (which included the off-rate selections step), strong enrichment factors in the range of 10-263 were determined.

**Table 1.**
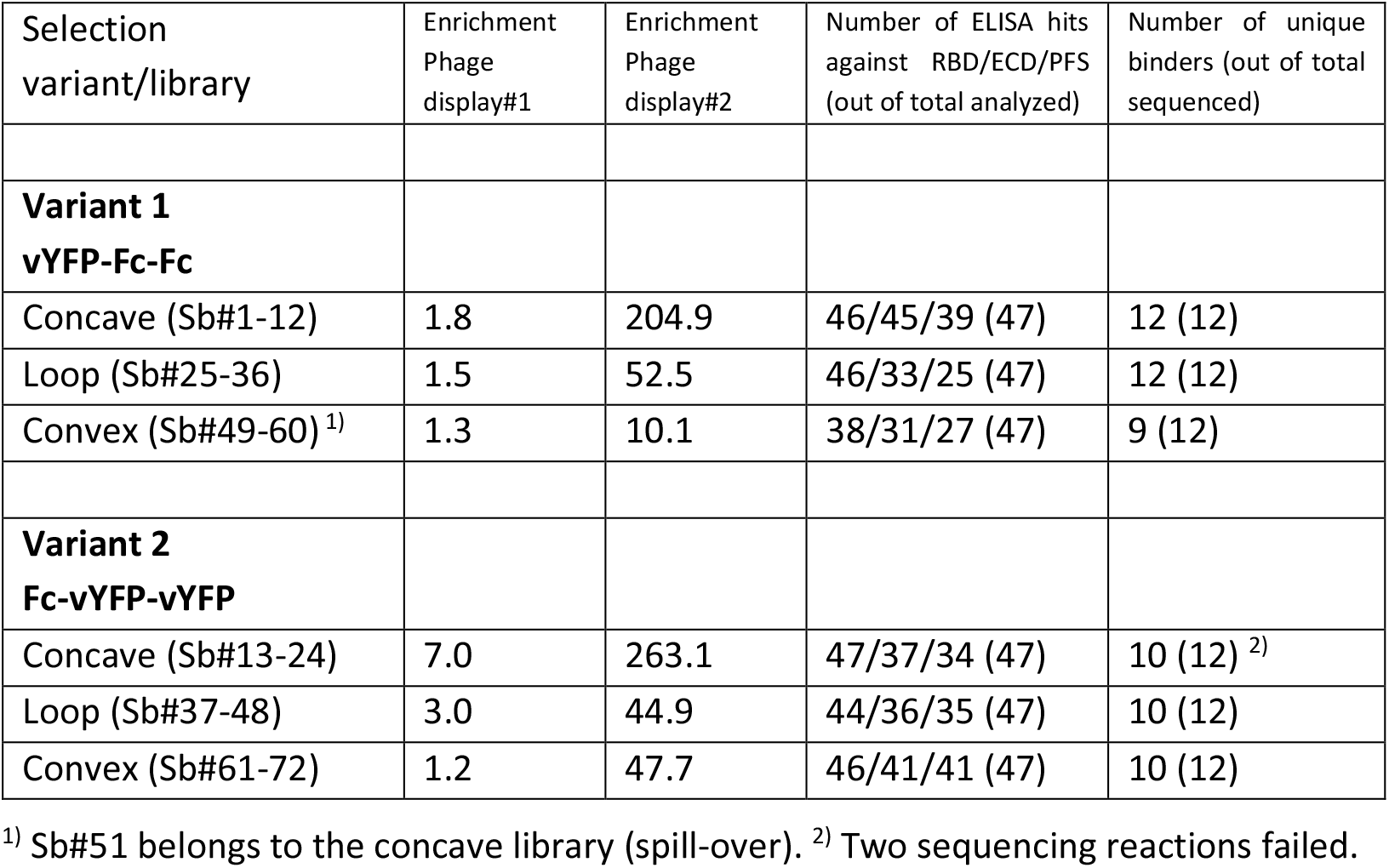
Key parameters of selection process

| Selection variant/library | Enrichment Phage display#1 | Enrichment Phage display#2 | Number of ELISA hits against RBD/ECD/PFS (out of total analyzed) | Number of unique binders (out of total sequenced) |
| --- | --- | --- | --- | --- |
| **Variant 1**<br>**vYFP-Fc-Fc** |  |  |  |  |
| Concave (Sb#1-12) | 1.8 | 204.9 | 46/45/39 (47) | 12 (12) |
| Loop (Sb#25-36) | 1.5 | 52.5 | 46/33/25 (47) | 12 (12) |
| Convex (Sb#49-60) <sup>1)</sup> | 1.3 | 10.1 | 38/31/27 (47) | 9 (12) |
| **Variant 2**<br>**Fc-vYFP-vYFP** |  |  |  |  |
| Concave (Sb#13-24) | 7.0 | 263.1 | 47/37/34 (47) | 10 (12) <sup>2)</sup> |
| Loop (Sb#37-48) | 3.0 | 44.9 | 44/36/35 (47) | 10 (12) |
| Convex (Sb#61-72) | 1.2 | 47.7 | 46/41/41 (47) | 10 (12) |

<sup>1)</sup> Sb#51 belongs to the concave library (spill-over). <sup>2)</sup> Two sequencing reactions failed.

### Sybody identification by ELISA

After sub-cloning the pools from the phage display vector pDX_init into the sybody expression vector pSb_init, 47 clones of each of the 6 selections reactions (Table 1, Fig. 1B) were picked at random and expressed in small scale. Our standard ELISA was initially performed using RBD-vYFP (RBD), spike ectodomain containing S1 and S2 (ECD), and maltose binding protein (MBP) as unrelated dummy protein. As outlined in the Materials and Methods section, ELISA analysis revealed very high hit rates for the RBD and the ECD ranging from 81 % to 100 % and 66 % to 96 %, respectively (Fig. 2, Table 1). The majority of the sybodies giving an ELISA signal to the RBD also gave a clear signal the full-length spike protein (Fig. 2). However, there was a total of 44 hits that only gave an ELISA signal for RBD-vYFP, but not for the ECD. This could be due to the presence of cryptic RBD epitopes that are not accessible in the context of the full-length spike protein, or the respective sybodies may recognize the vYFP portion of the RBD-vYFP construct, though the selection procedure clearly disfavors the latter explanation. Importantly, background binding to the dummy protein MBP was not observed for any of the analyzed sybodies, clearly showing that the binders are highly specific. We then sequenced 72 sybodies that were ELISA-positive against RBD-vYFP as well as the full-length spike (12 for each of the 6 selection reactions numbered from Sb#1-72, see also Fig. 1B).

**Figure 2.**
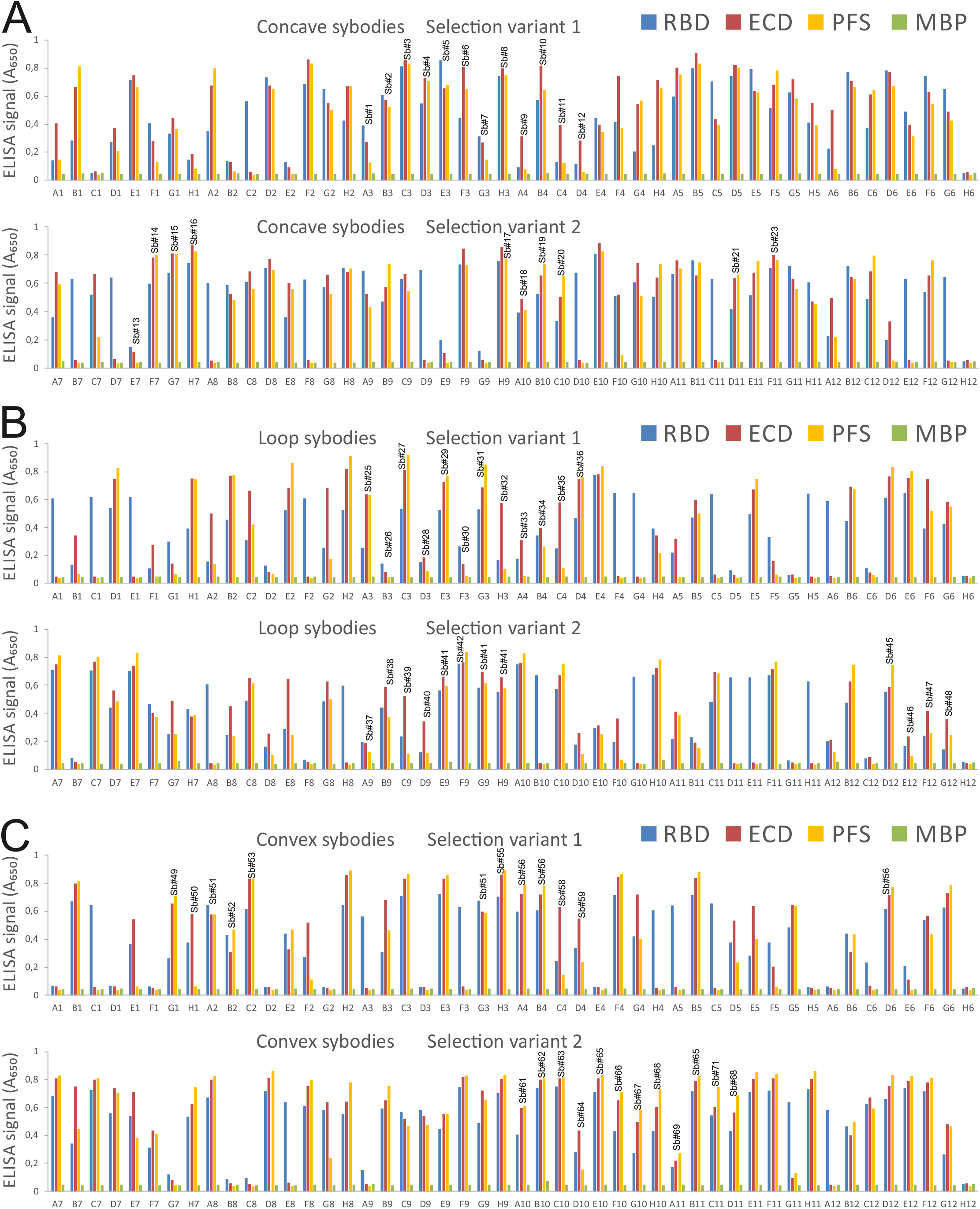
Sybody identification by ELISA. (**A**) Concave sybodies. (**B**) Loop sybodies. (**C**) Convex sybodies. For each of the six independent selection reactions, 47 clones were picked at random and analyzed by ELISA. A non-randomized sybody was used as negative control (wells H6 and H12, respectively). Sybodies that were sequenced are marked with the respective sybody name (Sb#1-72). Please note that identical sybodies that were found 2-3 times are marked with the same sybody name (e.g. Sb#41). ELISA analyses shown in these graphs were performed on three different days: (1) RBD and MBP, (2) ECD, (3) PFS.

Subsequent to sybody sequencing, we also performed the ELISA using engineered pre-fusion-stabilized spike ectodomain (PFS) (Fig. 2), which was not available at the onset of the project. Overall, the ELISA signals for the ECD and PFS are highly similar. However, there are around 40 sybodies that bind to the ECD clearly stronger than to the PFS (yet the opposite scenario was never observed). This could be explained by the fact that the PFS forms a trimer, while the oligomeric state of the ECD is not clear. In addition, the ECD might adopt partially or completely a post-fusion state, whereas PFS is expected to predominantly adopt the pre-fusion state. Trimer formation as well as pre-fusion stabilization might shield certain binding epitopes on the RBD in the context of the PFS, which might become accessible as the spike falls apart into monomers and/or transits to the post-fusion state. In light of our ELISA data, the PFS construct will be a crucial element in any future sybody selection campaigns.

### Sequence analysis

Sequencing results of 70 out of 72 sybody clones were unambiguous. Out of these 70 clones, 63 were found to be unique and the respective clone names are indicated in the ELISA figure (Fig. 2, Table 2). Of note, there were no duplicate binders identified in both selection variants, indicating that the two separate selection streams gave rise to completely different arrays of sybodies. As an additional note, one sybody identified from the supposed convex library turned out to belong to the concave library; spill-over of sybodies across libraries is occasionally observed. Hence, there was a total of 23 concave, 22 loop and 18 convex sybodies, which were then aligned according to their library origin (Figs. 3-5). As a final analysis, all sybody sequences were aligned to generate a phylogenetic tree, which shows a clear segregation across the three libraries and indicates a large sequence variability of the identified sybodies (Fig. 6).

**Table 2.**
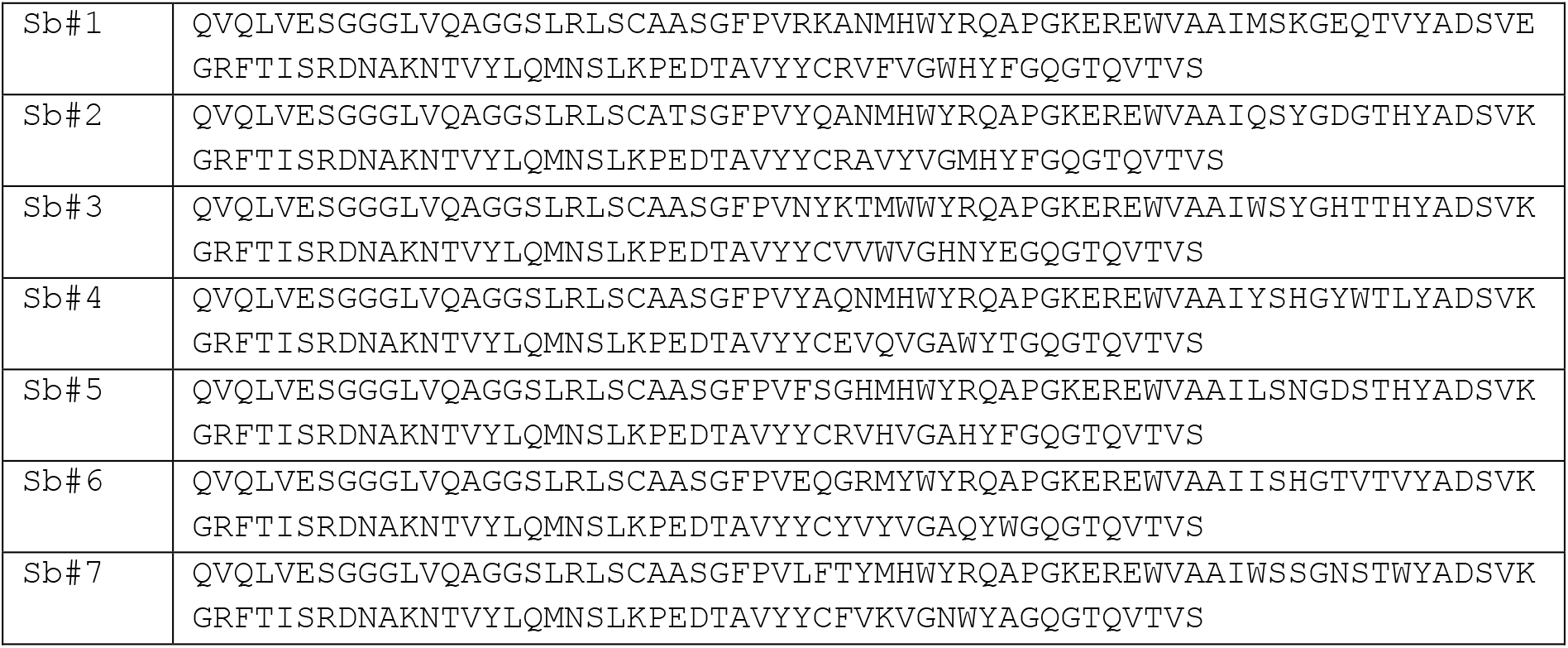
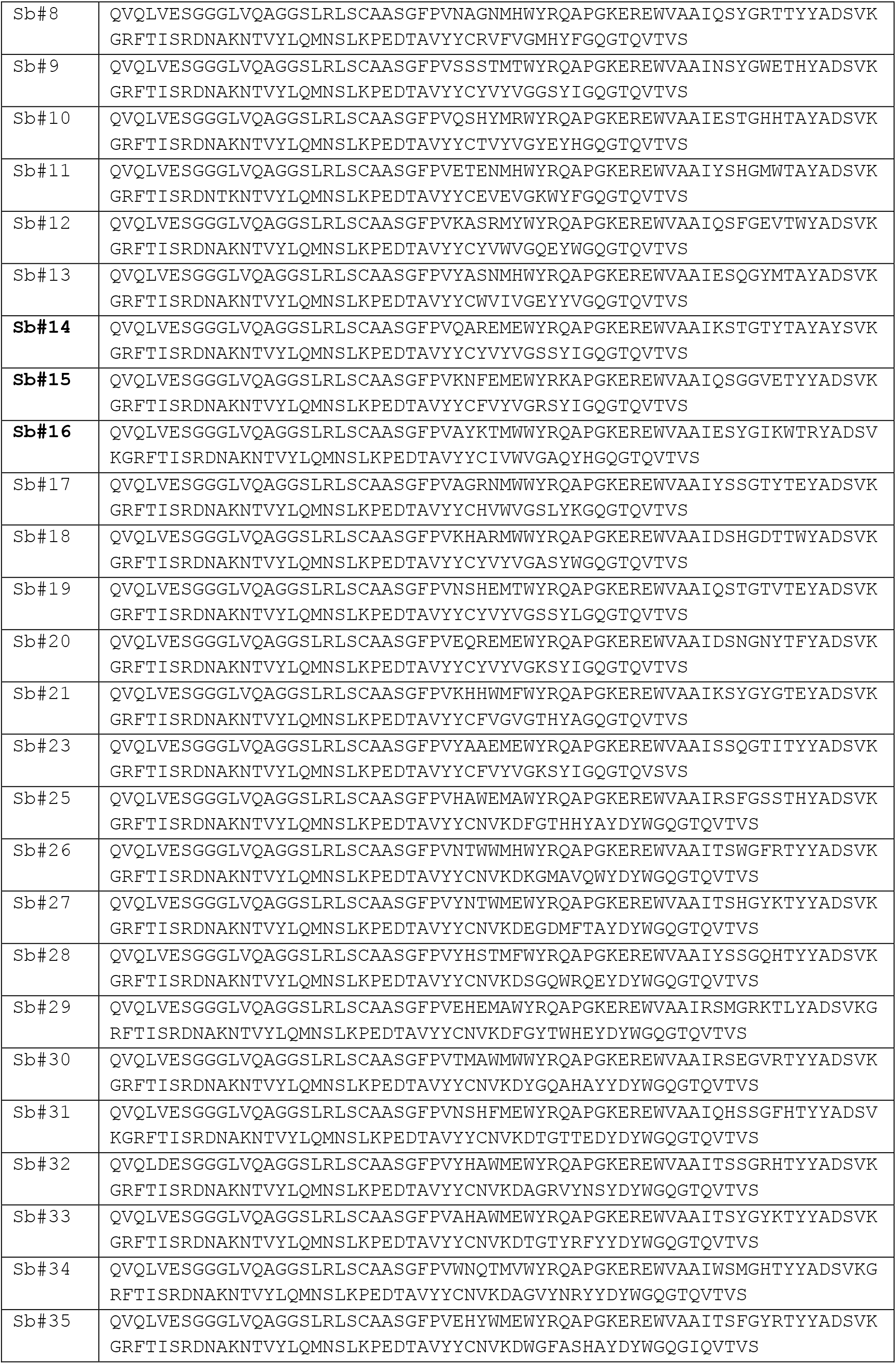
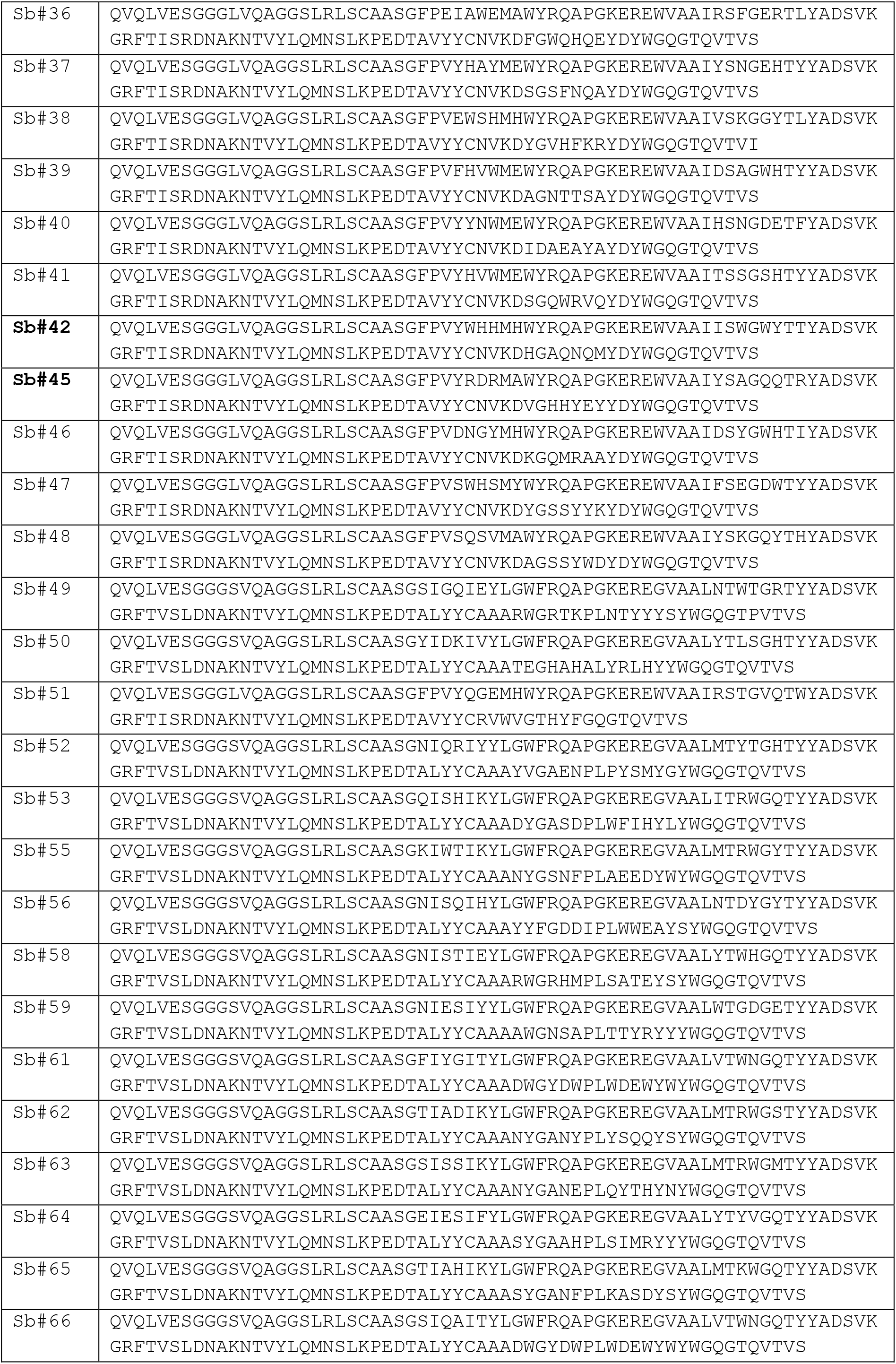
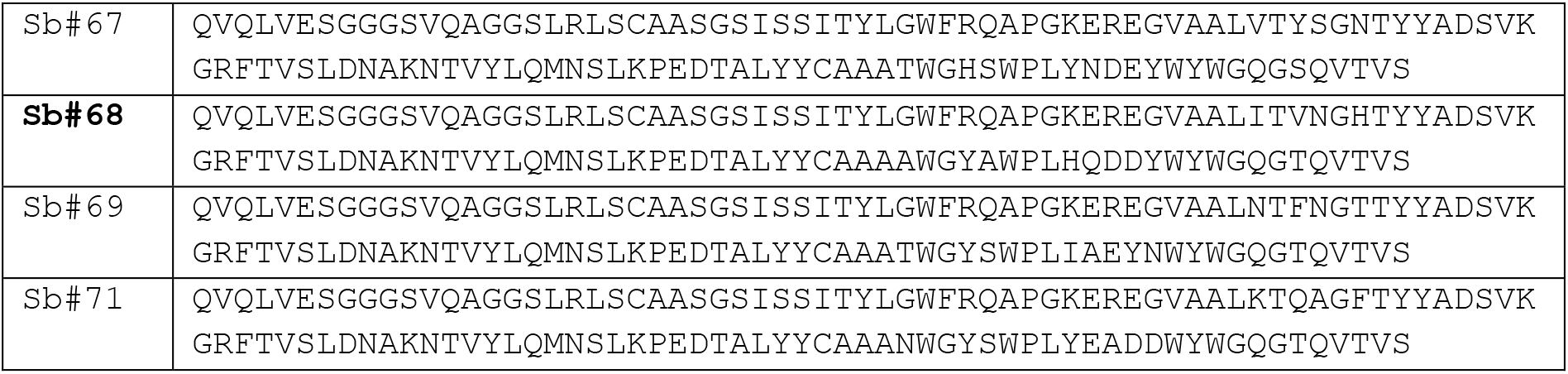
Sybody protein sequences

**Figure 3.**
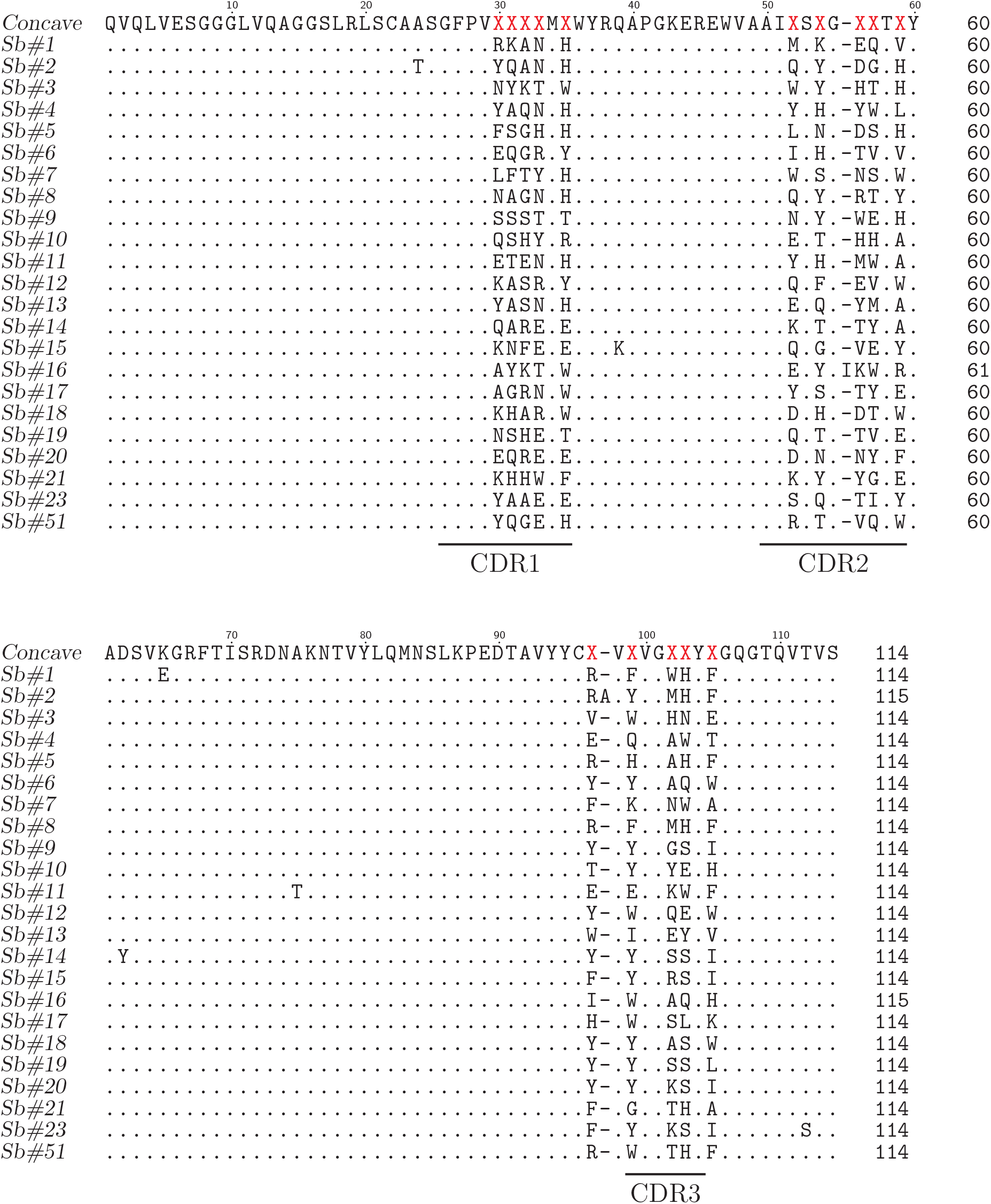
Sequence alignment of concave RBD sybodies.

**Figure 4.**
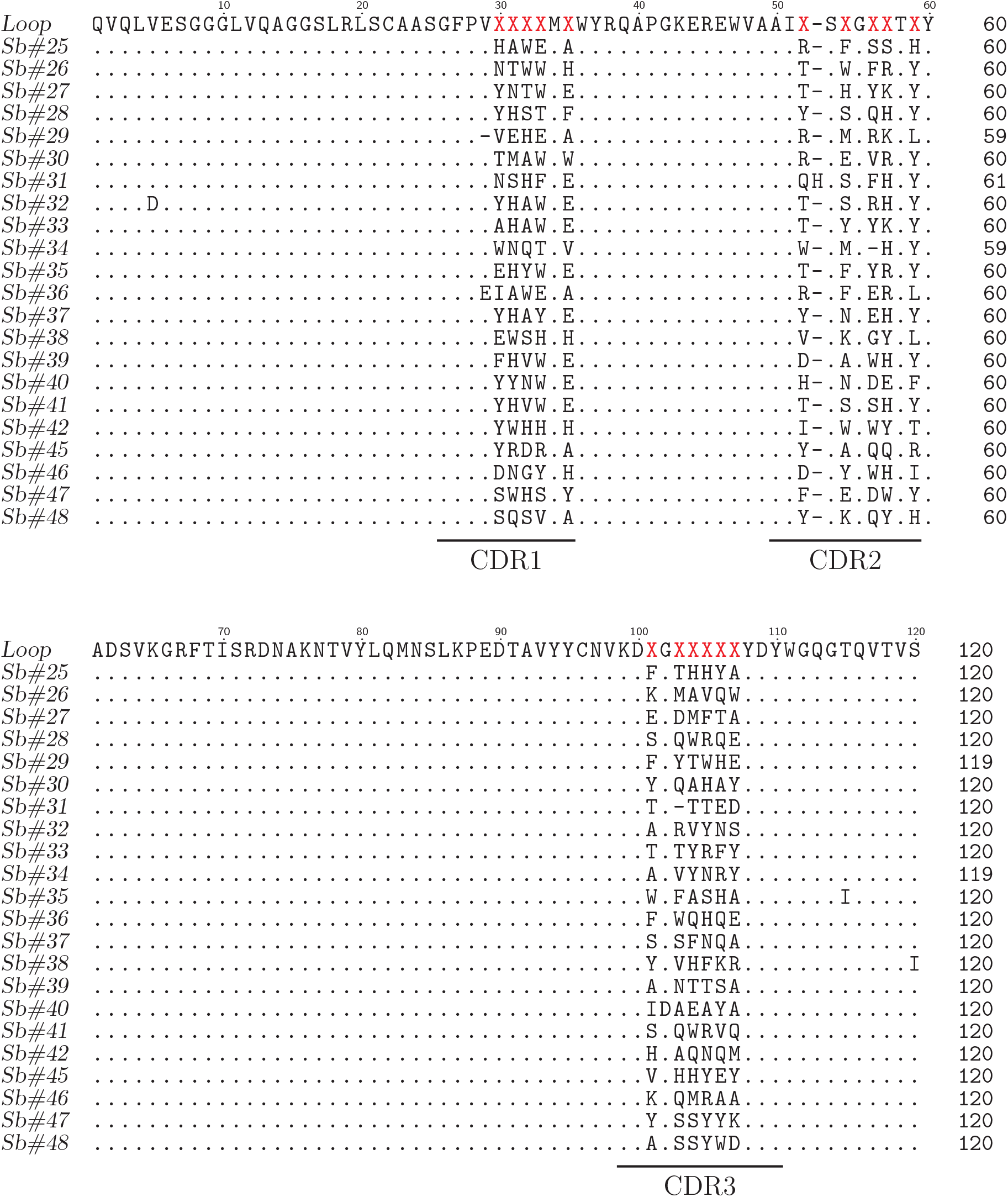
Sequence alignment of loop RBD sybodies.

**Figure 5.**
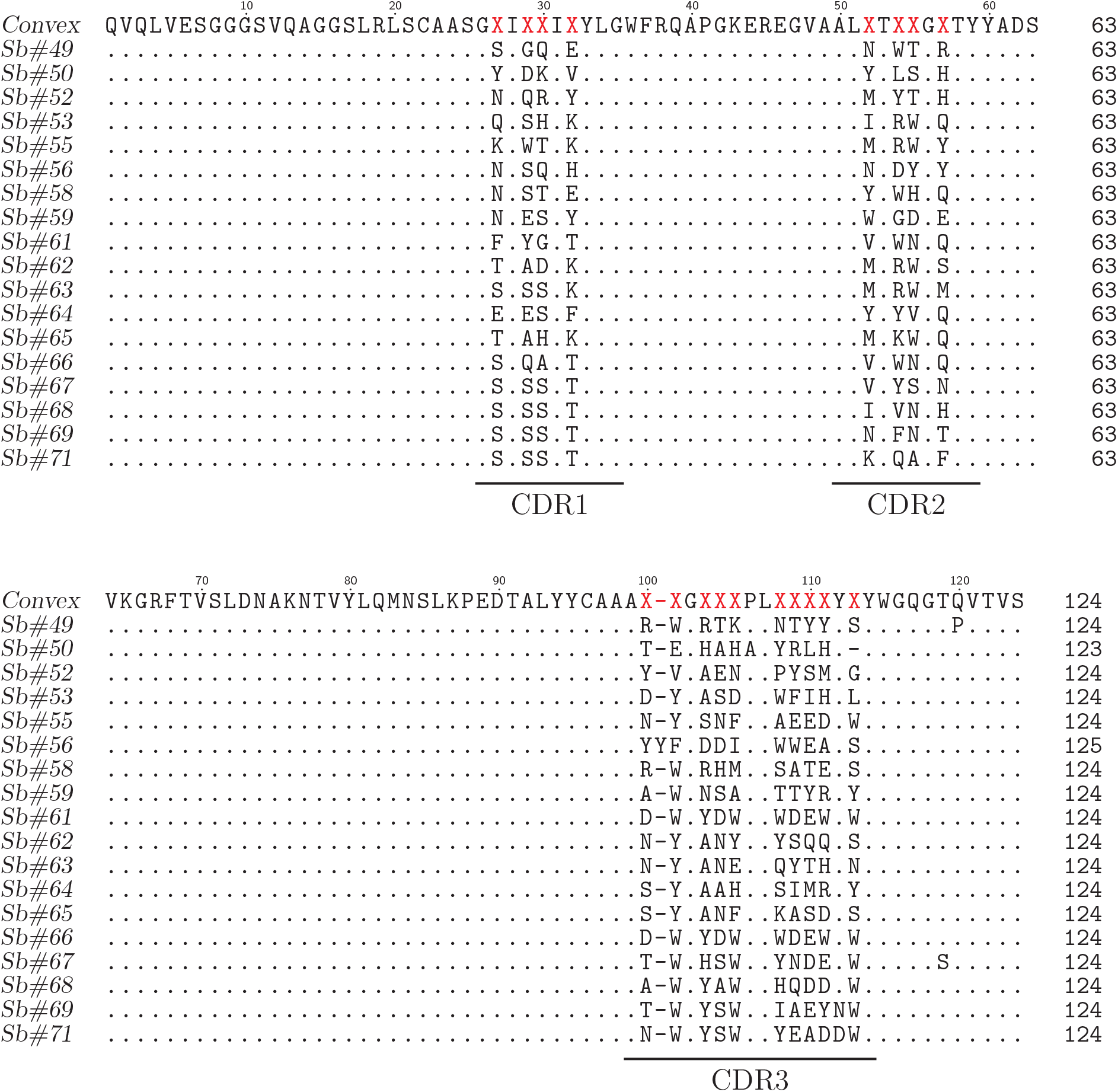
Sequence alignment of convex RBD sybodies.

**Figure 6.**
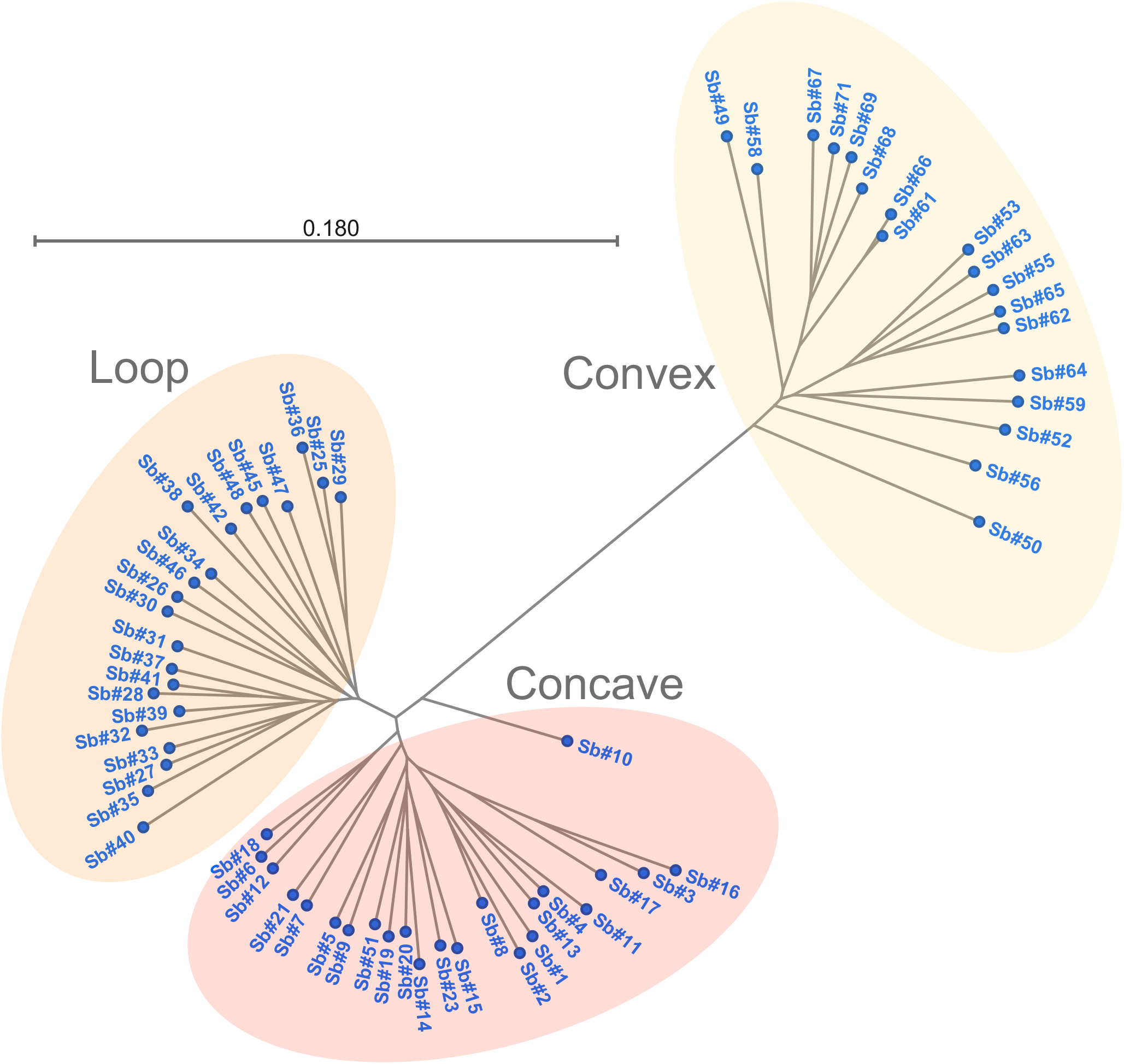
Phylogenetic tree of RBD sybodies. A radial tree was generated in CLC 8.1.3.

### Purification of sybodies and kinetic analysis of sybody interaction with SARS-CoV-2 proteins

The 63 unique sybodies were individually expressed in *E. coli* and purified via Ni-NTA affinity chromatography and gel filtration. Ultimately, 57 sybodies were sufficiently well-behaved, with respect to solubility, yield, and monodispersity, to proceed with further characterization. For a kinetic analysis of sybody interactions with the viral spike, we employed grating-coupled interferometry (GCI) to probe sybody binding to immobilized RBD-vYFP or ECD. First, the 57 purified sybodies were subjected to an off-rate screen, which revealed six sybodies (Sb#14, Sb#15, Sb#16, Sb#42, Sb#45, and Sb#68) with strong binding signals and comparatively slow off-rates. Binding constants were then determined by measuring on-and off-rates over a range of sybody concentrations, revealing affinities within a range of 20–180 nM to the SARS-CoV-2 spike (Fig. 7, Table 3). Of note, binding affinities were consistently equal or higher for the ECD as compared to the RBD-vYFP, in particular in case of Sb#68 for which the off-rate differs by more than two-fold. This might indicate a binding avidity effect arising from binding epitopes clustering in the context of the spike trimer or differences with regards to the glycan structures (RBD-vYFP was produced in HEK cells, whereas the ECD was produced in insect cells). To our surprise, the majority of purified and ELISA-positive sybodies (51 out of 57) displayed binding affinities worse than 200 nM. This may be attributed to the presence of complex heterogeneous Asn-linked glycans within the RBD, which could hinder the isolation of specific high-affinity binders. Alternatively, given that the final ELISA step of the selection process resulted in a substantial number of positive clones, insufficiently stringent conditions may have favored the high positive hit rate of low-affinity binders.

**Table 3.**
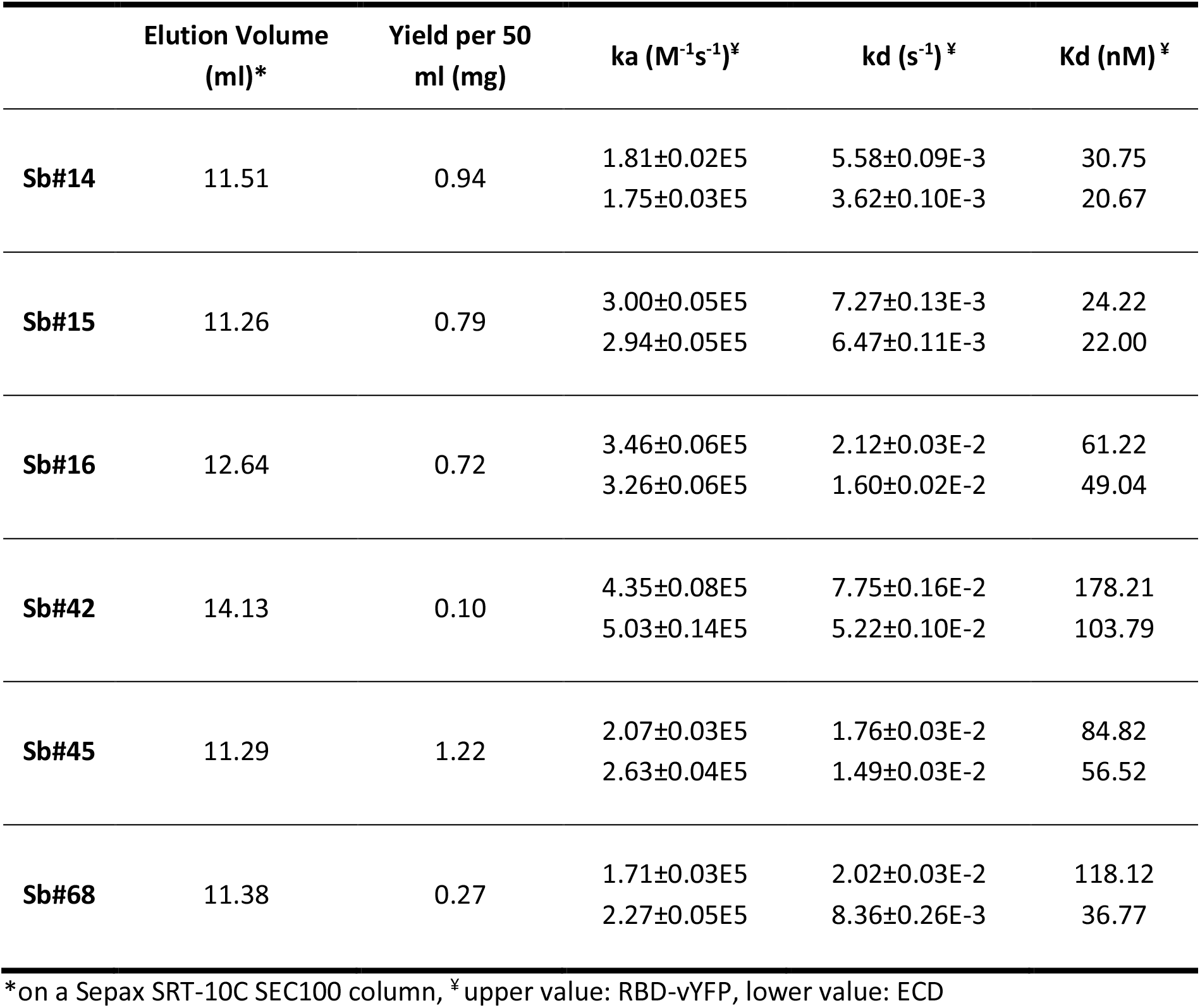
Purification details of top-performing sybodies, and kinetic parameters for sybody interactions with SARS-CoV-2 proteins

**Figure 7.**
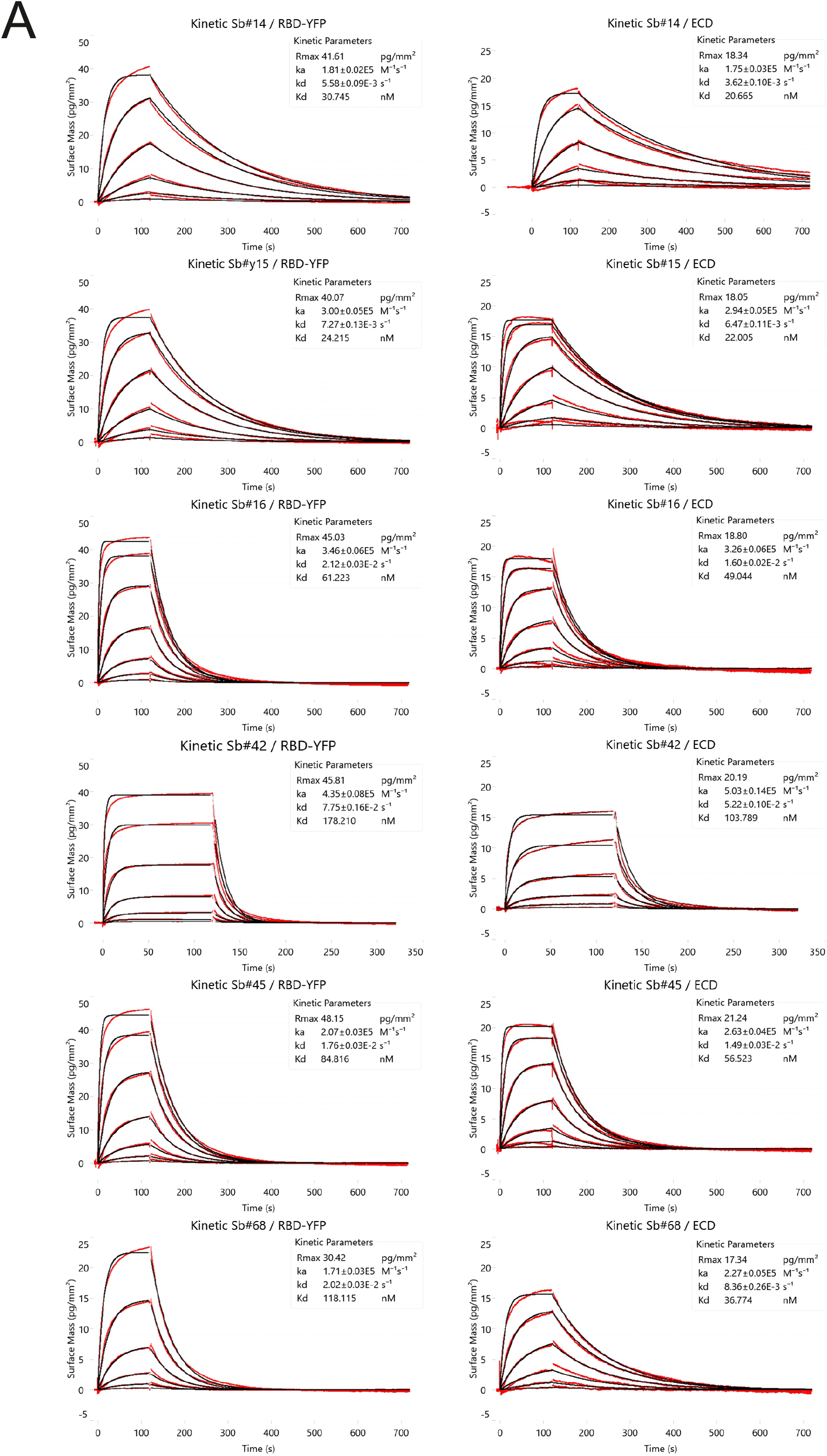
Kinetic characterization of the top six sybodies. (**A**) Binding kinetics were measured by grating-coupled interferometry on the WAVE system (Creoptix AG, Switzerland). RBD-vYFP and ECD were immobilized and the sybodies were injected at increasing concentrations ranging from 1.37 nM to 1 μM. Data were fitted using a Langmuir 1:1 model.

### ACE2 competition analysis

Since virulence of SARS-CoV-2 is dependent on the ability of the viral RBD to bind to human ACE2 (hACE2), we sought to determine which of the 57 selected sybodies that were well-behaved upon purification could inhibit interaction between the isolated RBD and purified hACE2. For this assessment, ELISA plates were coated with purified hACE2, and the binding of purified RBD to the immobilized hACE2 was measured in the presence or absence of an excess of each purified sybody (Fig. 8). While the absence of any added sybody resulted in a strong ELISA signal corresponding to RBD association with hACE2, the pre-incubation of nearly all sybodies with the RBD resulted in an attenuated signal, implying that these binders inhibit RBD-hACE2 association. This signal decrease relative to unchallenged RBD was modest for most sybodies, with an average signal reduction of about 50%, but five sybodies demonstrated exceptionally high apparent inhibition of RBD-hACE2 interaction (Sb#14, Sb#15, Sb#16, Sb#42, and Sb#45), showing ≥90% signal reduction. Notably, the aforementioned kinetic analysis had shown that these sybodies were also among the strongest RBD binders. Taken together, this data suggests that Sb#14, Sb#15, Sb#16, Sb#42, and Sb#45 recognize a surface region on the RBD that overlaps with the hACE2 binding site.

**Figure 8.**
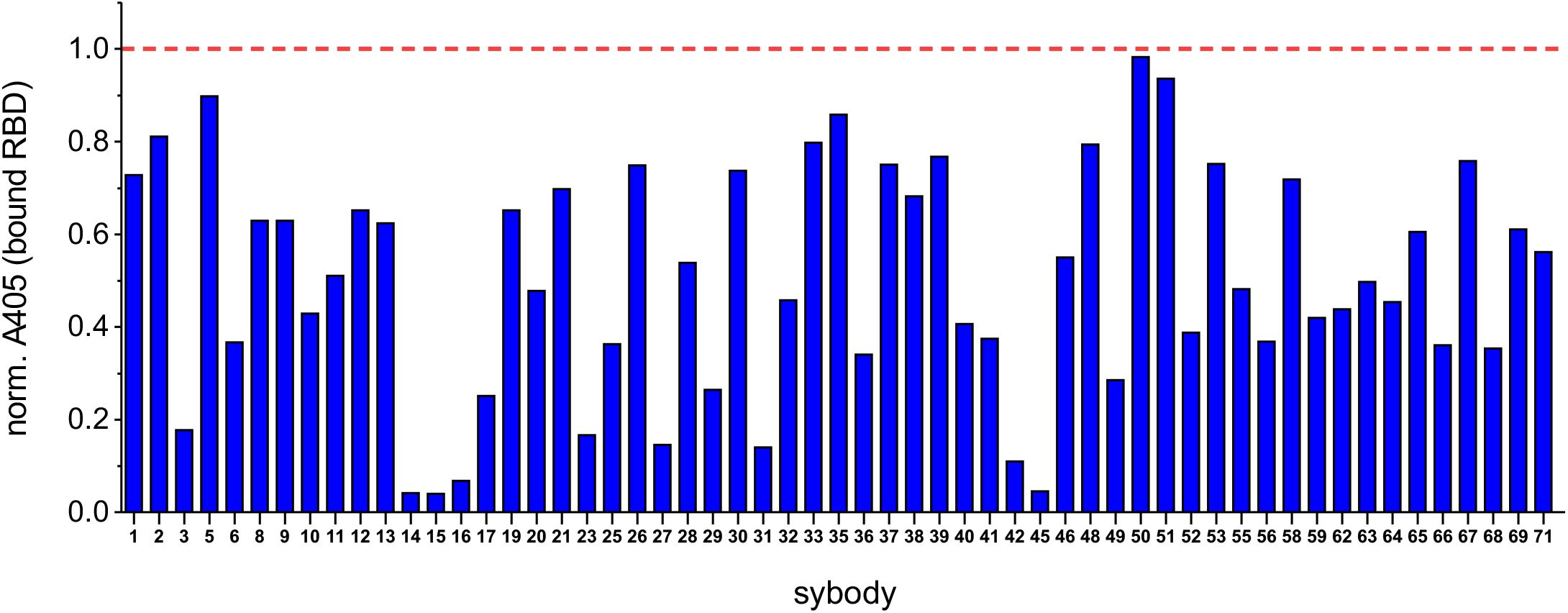
Sybodies inhibit RBD binding to ACE2. The effect of sybodies on RBD association with human ACE2 was assessed with an ELISA. Individual sybodies (500 nM, sybody number shown on X-axis) were incubated with biotinylated RBD-vYFP (25 nM) and the mixtures were exposed to immobilized ACE2. Bound RBD-vYFP was detected with streptavidin-peroxidase/TMB. Each column indicates background-subtracted absorbance at 405 nm, normalized to the signal corresponding to RBD-vYFP in the absence of sybody (dashed red line).

### Simultaneous binding of multiple sybodies to the RBD

While kinetic analysis had revealed Sb#68 to be among the stronger binders to the SARS-CoV-2 ectodomain (K_D_ ≈ 37 nM, Fig. 7, Table 3), the hACE2 competition ELISA revealed that Sb#68 does not inhibit hACE2-RBD interaction to the same extent as other sybodies with comparable affinities (65% inhibition for Sb#68, compared to >90% for Sb#14, Sb#15, Sb#16, and Sb#45). Therefore, it was hypothesized that Sb#68 may interact with a non- or partially-overlapping surface on the RBD, relative to the more strongly-inhibiting sybodies. Using Sb#15 as a representative of the hACE2-inhibiting sybodies, we analyzed the ability of Sb#15 and Sb#68 to simultaneously associate with the RBD. First, ELISA experiments demonstrate that incubation of Sb#68 with the pre-fusion spike only slightly prevents the spike from binding to immobilized Sb#15, whereas pre-incubation with Sb#14, Sb#15, Sb#16, Sb#42, or Sb#45 completely prevents spike interaction with immobilized Sb#15 (Fig. 9). In agreement with the ELISA data, GCI experiments revealed that co-injection of Sb#15 and Sb#68 results in a clear (but not fully additive) increase of the response signal, relative to Sb#15 or Sb#68 injected alone, implying simultaneous binding of Sb#15 and Sb#68 (Fig. 9). The control GCI experiment involving the co-injection of Sb#15 and Sb#45 did not result in a similar signal increase (Fig. 9). In sum, this data plausibly suggests that Sb#15 and Sb#68 can simultaneously bind to the RBD. For the design of therapeutics against SARS-CoV-2, the fusion of such a pair of non-overlapping binders could provide benefits via increased overall avidity to the spike protein.

**Figure 9.**
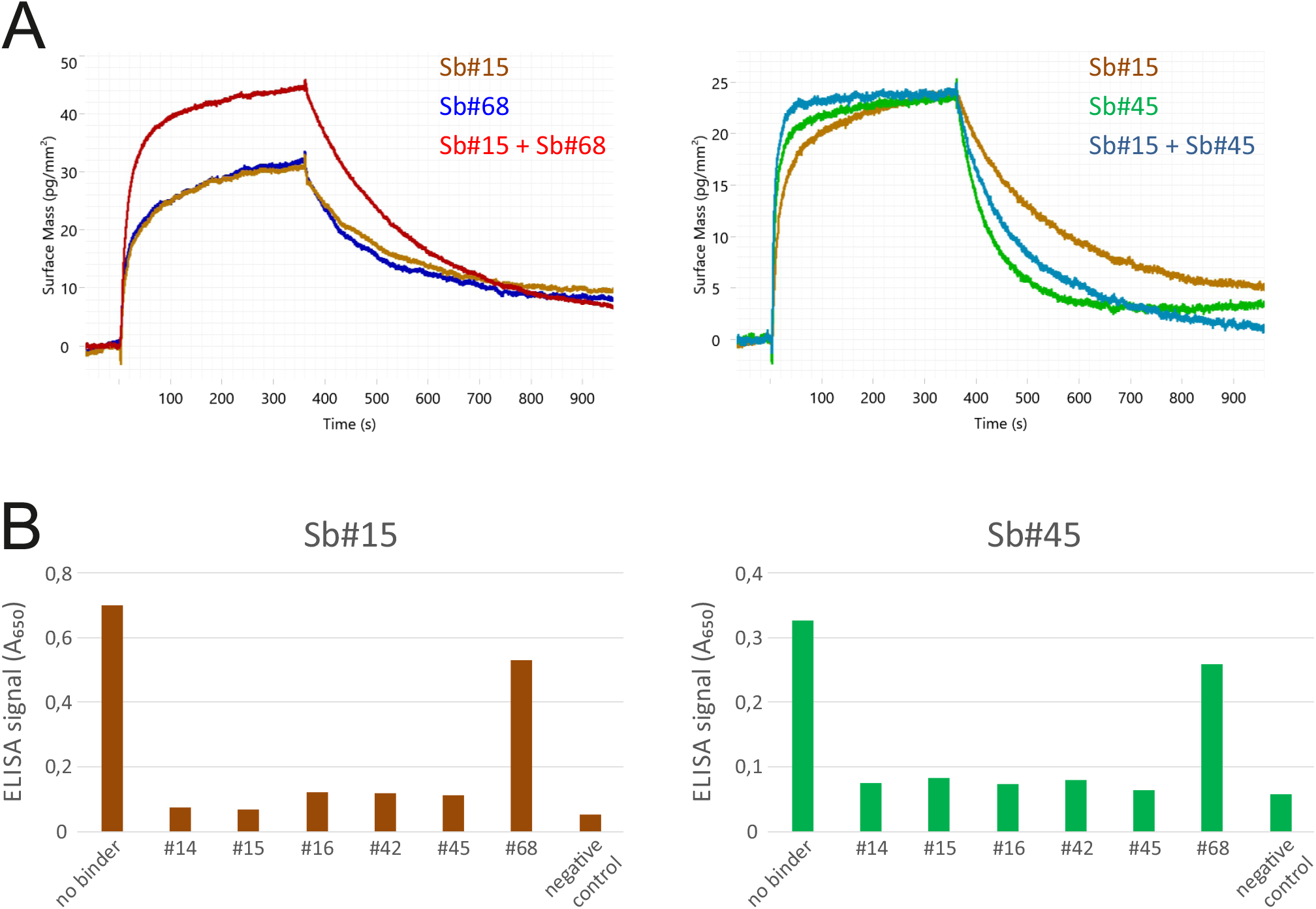
Simultaneous binding of Sb#15 and Sb#68. (**A**) Simultaneous binding of sybodies was analyzed using grating-coupled interferometry on the WAVE system (Creoptix AG, Switzerland). Biotinylated ECD was immobilized and the binders were injected alone and simultaneously at saturating concentrations (Sb#15: 200 nM, Sb#45: 500 nM, Sb#68: 500 nM). Superimposed sensorgrams are shown. (**B**) Competition ELISA. Title of the graphs indicate the sybody which was directly coated on the plate at a concentration of 25 nM. The labels on the x-axes depict the sybody used for competition. To determine the background signal, buffer devoid of protein was added.

### Conclusion and outlook

We have demonstrated the ability of our rapid *in vitro* selection platform to generate sybodies against the SARS-CoV-2 RBD, within a two-week timeframe. Characterization of these sybodies has identified a high-affinity subset of binders that also inhibit the RBD-ACE2 interaction. We anticipate that the presented panel of anti-RBD sybodies could be of use in the design of urgently required therapeutics to mitigate the COVID-19 pandemic, particularly in the development of inhalable prophylactic formulations [37]. Furthermore, our identification of a pair of sybodies that can simultaneously associate with the RBD may offer an attractive foundation for the construction of a polyvalent sybody-based therapeutic. We have attempted to provide a complete account of the generation of these molecules, including full sequences and detailed methods, such that other researchers may contribute to their ongoing analysis. Future work may include virus neutralization assays using the identified sybodies, as well as further selection campaigns targeting additional spike epitopes. Finally, our recently described flycode technology could be utilized for deeper interrogation of selection pools, in order to facilitate discovery of exceptional sybodies that possess very slow off-rates or recognize rare epitopes [41].

## METHODS

### Cloning, expression and purification of SARS-CoV-2 proteins

A gene encoding SARS-CoV-2 residues Pro330—Gly526 (RBD, GenBank accession QHD43416.1), downstream from a modified N-terminal human serum albumin secretion signal [42], was chemically synthesized (GeneUniversal). This gene was subcloned using FX technology [43] into a custom mammalian expression vector [44], appending a C-terminal 3C protease cleavage site, myc tag, Venus YFP[45], and streptavidin-binding peptide [46] onto the open reading frame (RBD-vYFP). 100–250 mL of suspension-adapted Expi293 cells (Thermo) were transiently transfected using Expifectamine according to the manufacturer protocol (Thermo), and expression was continued for 4–5 days in a humidified environment at 37°C, 8% CO_2_. Cells were pelleted (500*g*, 10 min), and culture supernatant was filtered (0.2 μm mesh size) before being passed three times over a gravity column containing NHS-agarose beads covalently coupled to the anti-GFP nanobody 3K1K [39], at a resin:culture ratio of 1ml resin per 100ml expression culture. Resin was washed with 20 column-volumes of RBD buffer (phosphate-buffered saline, pH 7.4, supplemented with additional 0.2M NaCl), and RBD-vYFP was eluted with 0.1 M glycine, pH 2.5, via sequential 0.5 ml fractions, without prolonged incubation of resin with the acidic elution buffer. Fractionation tubes were pre-filled with 1/10 vol 1M Tris, pH 9.0 (50 μl), such that elution fractions were immediately pH-neutralized. Fractions containing RBD-vYFP were pooled, concentrated, and stored at 4°C. Purity was estimated to be >95%, based on SDS-PAGE (not shown). Yield of RBD-vYFP was approximately 200–300 μg per 100 ml expression culture. A second purified RBD construct, consisting of SARS-CoV-2 residues Arg319—Phe541 fused to a murine IgG1 Fc domain (RBD-Fc) expressed in HEK293 cells, was purchased from Sino Biological (Catalogue number: 40592-V05H, 300 μg were ordered). Purified full-length spike ectodomain (ECD) comprising S1 and S2 (residues Val16—Pro1213) with a C-terminal His-tag and expressed in baculovirus-insect cells was purchased from Sino Biological (Catalogue number: 40589-V08B1, 700 μg were ordered). The prefusion ectodomain of the SARS-CoV2 Spike protein (residues 1-1208) [13], was transiently transfected into 50×10^8^ suspension-adapted ExpiCHO cells (Thermo Fisher) using 3 mg plasmid DNA and 15 mg of PEI MAX (Polysciences) per 1L ProCHO5 medium (Lonza) in a 3L Erlenmeyer flask (Corning) in an incubator shaker (Kühner). One hour post-transfection, dimethyl sulfoxide (DMSO; AppliChem) was added to 2% (v/v). Incubation with agitation was continued at 31°C for 5 days. 1L of filtered (0.22 um) cell culture supernatant was clarified. Then, a 1mL Gravity flow Strep-Tactin^®^XT Superflow^®^ column (iba lifescience) was rinsed with 2 ml buffer W (100 mM Tris, pH 8.0, 100 mM NaCl, 1 mM EDTA) using gravity flow. The supernatant was added to the column, which was then rinsed with 5 ml of buffer W (all with gravity flow). Finally, six elution steps were performed by adding each time 0.5 ml of buffer BXT (50mM Biotin in buffer W) to the resin. All purification steps were performed at 4°C.

### Biotinylation of target proteins

To remove amines, all proteins were first extensively dialyzed against RBD buffer. Proteins were concentrated to 25 μM using Amicon Ultra concentrator units with a molecular weight cutoff of 30 – 50 kDa. Subsequently, the proteins were chemically biotinylated for 30 min at 25°C using NHS-Biotin (Thermo Fisher, #20217) added at a 10-fold molar excess over target protein. Immediately after, the three samples were dialyzed against TBS pH 7.5. During these processes (first dialysis/ concentrating/ biotinylation/ second dialysis), 20 %, 30 %, 65 % and 44% of the RBD-vYFP, RBD-Fc, ECD and PFS respectively were lost due to sticking to the concentrator filter or due to aggregation. Biotinylated RBD-vYFP, RBD-Fc and ECD were diluted to 5 μM in TBS pH 7.5, 10 % glycerol and stored in small aliquots at −80°C. Biotinylated PFS was stored at 4°C in TBS pH 7.5.

### Sybody selections

Sybody selections with the three sybody libraries concave, loop and convex were carried out as described in detail before [36]. In short, one round of ribosome display followed by two rounds of phage display were carried out. Binders were selected against two different constructs of the SARS-CoV-2 RBD; an RBD-vYFP fusion and an RBD-Fc fusion. MBP was used as background control to determine the enrichment score by qPCR [36]. In order to avoid enrichment of binders against the fusion proteins (YFP and Fc), we switched the two targets after ribosome display (Fig. 1B). For the off-rate selections we did not use non-biotinylated target proteins as described in the standard protocol, because we did not have enough purified protein at hand to do so. Instead we sub-cloned all three libraries for both selections after the first round of phage display into the pSb_init vector (10^8^ clones) and expressed the six pools in *E. coli* MC1061 cells. Then the pools corresponding to the same selection were pooled for purification. The two final pools were purified by Ni-NTA resin using gravity flow columns, followed by buffer exchange of the main peak fraction using a desalting PD10 column in TBS pH 7.5 to remove imidazole. The pools were eluted with 3.2 ml instead of 3.5 ml TBS pH 7.5 in order to ensure complete buffer exchange. These two purified pools were used for the off-rate selection in the second round of phage display at concentrations of approximately 390 μM for selection variant 1 (RBP-Fc) and 450 μM for selection variant 2 (RBP-YFP). The volume used for off-rate selection was 500 μl. Just before the pools were used for the off-rate selection, 0.5% BSA and 0.05% Tween-20 was added to each sample. Off-rate selections were performed for 3 minutes.

### Sybody identification by ELISA

ELISAs were performed as described in detail before [36]. 47 single clones were analyzed for each library of each selection. Since the RBD-Fc construct was incompatible with our ELISA format due to the inclusion of Protein A to capture an α-myc antibody, ELISA was performed only for the RBD-vYFP (50 nM) and the ECD (25 nM) and later on with the PFS (25 nM). Of note, the three targets were analyzed in three separate ELISAs. As negative control to assess background binding of sybodies, we used biotinylated MBP (50 nM). 72 positive ELISA hits were sequenced (Microsynth, Switzerland).

### Expression and Purification of sybodies

The 63 unique sybodies were expressed and purified as described [36]. In short, all 63 sybodies were expressed overnight in *E.coli* MC1061 cells in 50 ml cultures. The next day the sybodies were extracted from the periplasm and purified by Ni-NTA affinity chromatography (batch binding) followed by size-exclusion chromatography using a Sepax SRT-10C SEC100 size-exclusion chromatography (SEC) column equilibrated in TBS, pH 7.5, containing 0.05% (v/v) Tween-20 (detergent was added for subsequent kinetic measurements). Six out of the 63 binders (Sb#4, Sb#7, Sb#18, Sb#34, Sb#47, Sb#61) were excluded from further analysis due to suboptimal behavior during SEC analysis (i.e. aggregation or excessive column matrix interaction).

### Grating-coupled interferometry (GCI)

Kinetic characterization of sybodies binding onto SARS-CoV-2 spike proteins was performed using GCI on the WAVEsystem (Creoptix AG, Switzerland), a label-free biosensor. Biotinylated RBD-vYFP and ECD were captured onto a Streptavidin PCP-STA WAVEchip (polycarboxylate quasi-planar surface; Creoptix AG) to a density of 1300-1800 pg/mm^2^. Sybodies were first analyzed by an off-rate screen performed at a concentration of 200 nM (data not shown) to identify binders with sufficiently high affinities. The six sybodies Sb#14, Sb#15, Sb#16, Sb#42, Sb#45, and Sb#68 were then injected at increasing concentrations ranging from 1.37 nM to 1 μM (three-fold serial dilution, 7 concentrations) in TBS buffer supplemented with 0.05 % Tween-20. Sybodies were injected for 120 s at a flow rate of 30 μl/min per channel and dissociation was set to 600 s to allow the return to baseline. Sensorgrams were recorded at 25 °C and the data analyzed on the WAVEcontrol (Creoptix AG). Data were double-referenced by subtracting the signals from blank injections and from the reference channel. A Langmuir 1:1 model was used for data fitting.

### ACE2 competition ELISA

Purified recombinant hACE2 protein (MyBioSource, Cat# MBS8248492) was diluted to 10 nM in phosphate-buffered saline (PBS), pH 7.4, and 100 μl aliquots were incubated overnight on Nunc MaxiSorp 96-well ELISA plates (ThermoFisher #44-2404-21) at 4°C. ELISA plates were washed three times with 250 μl TBS containing 0.05% (v/v) Tween-20 (TBST). Plates were blocked with 250 μl of 0.5% (w/v) BSA in TBS for 2 h at room temperature. 100 μl samples of biotinylated RBD-vYFP (25 nM) mixed with individual purified sybodies (500 nM) were prepared in TBS containing 0.5% (w/v) BSA and 0.05% (v/v) Tween-20 (TBS-BSA-T) and incubated for 1.5 h at room temperature. These 100 μl RBD-sybody mixtures were transferred to the plate and incubated for 30 minutes at room temperature. 100 μl of streptavidin-peroxidase (Merck, Cat#S2438) diluted 1:5000 in TBS-BSA-T was incubated on the plate for 1 h. Finally, to detect bound biotinylated RBD-vYFP, 100 μl of development reagent containing 3,3′,5,5′-Tetramethylbenzidine (TMB), prepared as previously described [36], was added, color development was quenched after 3-5 min via addition of 100 μl 0.2 M sulfuric acid, and absorbance at 405 nm was measured. Background-subtracted absorbance values were normalized to the signal corresponding to RBD-vYFP in the absence of added sybodies.

### Dual-sybody competition ELISA

Purified sybodies carrying a C-terminal myc-His Tag (Sb_init expression vector) were diluted to 25 nM in 100 μl PBS pH 7.4 and directly coated on Nunc MaxiSorp 96-well plates (ThermoFisher #44-2404-21) at 4°C overnight. The plates were washed once with 250 μl TBS pH 7.5 per well followed by blocking with 250 μl TBS pH 7.5 containing 0.5% (w/v) BSA per well. In parallel, chemically biotinylated prefusion Spike protein (PFS) at a concentration of 10 nM was incubated with 500 nM sybodies for 1 h at room temperature in TBS-BSA-T. The plates were washed three times with 250 μl TBS-T per well. Then, 100 μl of the PFS-sybody mixtures were added to the corresponding wells and incubated for 3 min, followed by washing three times with 250 μl TBS-T per well. 100 μl Streptavidin-peroxidase polymer (Merck, Cat#S2438) diluted 1:5000 in TBS-BSA-T was added to each well and incubated for 10 min, followed by washing three times with 250 μl TBS-T per well. Finally, to detect PFS bound to the immobilized sybodies, 100 μl ELISA developing buffer (prepared as described previously [36]) was added to each well, incubated for 1 h (due to low signal) and absorbance was measured at 650 nm. As a negative control, TBS-BSA-T devoid of protein was added to the corresponding wells instead of a PFS-sybody mixture.

## DATA AVAILABILITY STATEMENT

The plasmids encoding for the six highest affinity binders will very soon be available through Addgene (Addgene #153522 - #153527).

## ACKNOWLEDGEMENTS

We thank Rony Nehmé and André Heuer (Creoptix AG, Wädeswil, Switzerland) for the acquisition, fitting and interpretation of GCI measurements using the WAVEsystem. We thank Florence Projer, David Hacker and Kelvin Lau (Protein Production and Structure Core Facility, EPFL, Switzerland) for the production of the pre-fusion spike protein. We are grateful to Jason McLellan (The University of Texas at Austin, U.S.) for having provided the pre-fusion-stabilized soluble spike expression vector.

